# *Chameleon* microRNAs in breast cancer: their elusive role as regulatory factors in cancer progression

**DOI:** 10.1101/2020.12.15.422846

**Authors:** Cesare Miglioli, Gaetan Bakalli, Samuel Orso, Mucyo Karemera, Roberto Molinari, Stéphane Guerrier, Nabil Mili

## Abstract

Breast cancer is one of the most frequent cancers affecting women. Non-coding micro RNAs (miRNAs) seem to play an important role in the regulation of pathways involved in tumor occurrence and progression. Extending on the research in Haakensen *et al*., where significant miRNAs were selected as being associated with the progression from normal breast tissue to breast cancer, in this work we put forward 112 sets of miRNA combinations, each including at most 5 expressions with high accuracy in discriminating healthy breast tissue from breast carcinoma. Our results are based on a recently developed machine learning technique which, instead of selecting a single model (or combination of features), delivers a set of models with equivalent predictive capabilities that allow to interpret and visualize the interaction of these features. These results shed new light on the biological action of the selected miRNAs which can behave in different ways according to the miRNA network with which they interact.

Indeed, these revealed connections may contribute to explain why, in some cases, different studies attribute opposite functions to the same miRNA. It is therefore possible to understand how the role of a genomic variable may change when considered in interaction with other sets of variables, as opposed to only considering its effect when it is evaluated within a unique combination of features. The approach proposed in this work provides a statistical basis for the notion of *chameleon* miRNAs and is inspired by the emerging field of systems biology.

**Author Summary:**

- The notion of a single predictive genomic (statistical) model is replaced by that of a set of models that can be considered as exchangeable due to their indistinguishable (optimal) predictive abilities;
- Our results indicate that the role of miRNAs cannot be interpreted independently from the combination of features with which they interact and can therefore vary considerably when considered in a network of different combinations. Some miRNAs may act as *chameleons* and behave in opposite manners thereby showing some kind of antagonistic duality;
- Some miRNAs are exchangeable inside models with equivalent predictive ability and seem to point to latent biological functions.

## Introduction

### Non-coding RNA and breast cancer

Breast cancer (BC) is the second-most common cancer and second-leading cause of cancer mortality in American women. In the USA, its incidence in 2019 was roughly 268,600 and it is responsible for an estimated 41,760 deaths [2] to which one must add 62,930 new cases of Ductal Carcinoma *In Situ* (DCIS). Dysregulation of microRNAs (miRNAs) plays a key role in almost all cancers, including BC [3]. miRNAs are short endogenous noncoding RNAs that regulate their target messenger RNAs (mRNA) by promoting mRNA degradation or repressing translation. Chang *et al*. [3] found that increased expression of 12 mature miRNAs — `hsa-miR-320a`, `hsa-miR-361-5p`, `hsa-miR-103a-3p`, `hsa-miR-21-5p`, `hsa-miR-374b-5p`, `hsa-miR-140-3p`, `hsa-miR-25-3p`, `hsa-miR-651-5p`,`hsa-miR-200c-3p`, `hsa-miR-30a-5p`, `hsa-miR-30c-5p`, and `hsa-let-7i-5p` — all predicted improved BC survival. In a recent review, Adhami *et al*. [4] determined that two miRNAs (`hsa-miR-21` and `hsa-miR-210`) were upregulated consistently and six miRNAs (`hsa-miR-145, hsa-miR-139-5p, hsa-miR-195, hsa-miR-99a, hsa-miR-497 and hsa-miR-205`) were downregulated in at least three studies. In another study, Haakensen *et al*. [1] identified some miRNA alterations during BC progression. These alterations were involved in the invasive signatures of BC including downregulation of `hsa-miR-139-5p` in aggressive subtypes and upregulation of `hsa-miR-29c-5p` in luminal subtypes. A total of 27 miRNAs were implicated in their proposed DCIS signature.

The latter study provided one of the main reasons to develop the work presented here. Indeed, Haakensen *et al*. [1] provide the following statement in their work:

“*hsa-miRNA-210-3p was significantly upregulated in both our analyses, but was downregulated in the same transition in Volinia et al* [13] *and is hence excluded from our proposed signature*”.In the wake of this statement, `hsa-miR-210` has previously been identified as a marker of poor prognosis in BC and other carcinomas [7]. In fact, Volinia *et al*. [13] found `hsa-miR-210` to be downregulated in DCIS compared with normal breast tissue, but upregulated in invasive carcinomas compared with DCIS. In addition, Shao *et al*. [23] recently showed that `hsa-miR-210` is associated with internal organ metastasis (liver, lung, and brain) in BC. In Haakensen’s study, `hsa-miR-210` was upregulated in DCIS compared to normal tissue and was not detected as significantly altered in any invasive subtype. Given the unclear role of this miRNA in breast carcinogenesis, Haakensen’s study therefore discarded it from the list of miRNAs involved in BC progression.

Considering these studies, this work aims at addressing some questions that naturally arise from their conclusions. The first of these questions is as follows: why weren’t there any concordant findings in many research studies regarding the role of miRNAs in the progression of BC? Are the different outcomes due to population selection, batch effect or to biological causes such as disease heterogeneity, overlapping of miRNA functions or network effects? A second question, that stems from the latter points, is the following: could an miRNA have either an activating effect or an inhibiting one in a given biological process (such as cancer progression) according to the other miRNAs with which it interacts? In other words, could a specific miRNA be upregulated in one study and downregulated in another as a result of the complex pathways in which it is involved (instead of this being the effect of the experimental conditions)? This work aims at investigating these questions more thoroughly and, for this purpose, we analyze the AHUS dataset using a recently proposed machine learning algorithm. The dataset is made available by Haakensen *et al*. on the open access ArrayExpress platform at: https://www.ebi.ac.uk/arrayexpress/experiments/E-MTAB-3759/?query=AHUS.

### Genomic connections in systems biology

The use of mathematical and computational models in biology is referred to as systems biology (SB) [21]. SB is a field which, among others, focuses on the assumption that a discrete biological function can rarely be attributed to a single molecule [20]. Instead, most biological characteristics arise from complex interactions among the cell’s numerous constituents, such as proteins, DNA, RNA and small molecules. Understanding the structure and the dynamics of complex intercellular networks that contribute to the structure and function of a living cell is therefore mandatory before assigning a function to any biological feature [19]. According to Barabási *et al*. [18], the inter- and intra-cellular connectivity implies that the impact of a specific genetic abnormality is not restricted to the activity of the gene product that carries it, but can spread along the links of the network and alter the activity of gene products that otherwise carry no defects. However the biological networks in which a single genomic variable is involved remain unknown. As a first step, one should then rely on a data-driven network built using statistical (and not biological) associations.

### Research questions

Our study has three goals: (i) to investigate if, and to what level of accuracy, it is possible to use different combinations of miRNAs as biomarkers to discriminate normal breast tissue from breast carcinoma; (ii) to check how the behavior of these miRNAs varies according to the specific combination with which it interacts; (iii) to search for interchangeable miRNAs in these predictive models and by doing so, to decipher the biological targets of these variables.

## Materials and Methods

### Genomic study

The results of our research are based on the AHUS dataset made available on the ArrayExpress platform by Haakensen *et al*. [1]. According to the authors, in order to collect this data the Akershus University Hospital sequentially collected breast tissue specimens from BC patients and from women undergoing surgery for breast reduction. These specimens were collected over 7 years between 2003 and 2009. miRNA-expression profiling was obtained for 55 invasive carcinomas and 70 normal breast tissue samples (including 29 tumor-adjacent normal tissue samples and 41 breast reduction samples) for a total of 125 as stated on the ArrayExpress platform. The samples were hybridized on Agilent 8×15K arrays (Agilent Technologies, Santa Clara, CA), catalogue number 4470B (v2) and 4470C (v3), and the features were extracted using Agilent Feature Extraction. Relevant information can be found in Haakensen *et al*. [1].

### Statistical analysis

When considering the research goals defined earlier, the statistical tools used to achieve them need to be defined accordingly. Hence, the first step is to find “different combinations of miRNAs” which implies that we are not aiming to find a single statistical (machine learning) model to classify normal breast tissue and breast carcinoma. Indeed, we intend to find a variety of models (miRNA combinations) that all perform this classification task with a high level of accuracy and renders them equivalent in terms of predictive power. The idea of considering a multitude of models is not a common one but has been put forward in different settings (see e.g. Caruana *et al*. [55]) and was adequately stressed, for example, in Whittingham *et al*. [56] who state that “[…*] further analysis should not be based on a single best model, but should explicitly acknowledge uncertainty among models that are similarly consistent with the data*”. In fact, depending on the setting, the reliance on a single model can be rather risky and can often deliver contradicting results regarding if and how certain variables contribute to explain or predict a given phenomenon of interest. In this perspective, we should choose an approach that allows to find a variety of “strong” models and that, in accordance with the subsequent research goals of this work, can be used to create miRNA networks highlighting how, for example, a specific miRNA can be used to detect (and can contribute differently to) breast carcinoma when considered with other miRNA combinations. In addition, in order to create networks that can also be interpreted from a biological perspective, we need these models also to be based on small (sparse) combinations of miRNAs.

There exist a wide variety of statistical and machine learning approaches to select and estimate models with few features (miRNAs) but, in most cases, these only select one model which therefore limits the possibility of considering how the impact of an miRNA can change when considered with another set of variables. For this reason this work proposes to use the “Sparse Wrapper AlGorithm” (*SWAG*) put forward in Molinari *et al*. [57] which is described in the following paragraphs.

### A wrapper method for sparse learning

The *SWAG* is a method derived from the Panning algorithm presented in Guerrier *et al*. [9] for gene selection problems. The premise of this method is the assumption that, in order to adequately predict a certain outcome of interest (e.g. breast carcinoma), we only need an extremely small set of features and that there are many models (combinations of small sets of features) that can all have equivalent and high predictive power. Aside from allowing to understand if and how certain features can behave differently when considered in presence with other sets of features, the output of the *SWAG* also allows to facilitate replicability of results. Indeed, when a study proposes a single model (and hence a single combination of features) in order to detect or predict a certain response, this may not be usable for a research or medical structure that may not have the possibility of measuring all the selected features. In order to respond to the above needs, the *SWAG* consists in a “greedy” wrapper algorithm that firstly requires the user to specify a model (or learning method), such as a logistic model, as well as the maximum number of variables (*p*_*max*_) to be considered within such a model. The latter choice can be made, for example, based on prior knowledge of the problem and interpretability requirements (the smaller this number, the easier the output will be interpreted). Based on these choices and supposing there is a total of *p* features (e.g. biomarkers), the *SWAG* starts through a first screening step where *p* models are built, each using a distinct feature. At this stage, the out-of-sample prediction error of each model can be estimated via *k*-fold cross-validation (CV) repeated *m* times and the best of these models (in terms of lowest prediction error) can be selected thereby providing a list of features that, on their own, appear to be highly predictive for the considered response. The definition of “best” models will be given by the user through a parameter *α* which represents a proportion (or percentile) and is usually chosen to be considerably small (i.e. between 0.01 and 0.1). With smaller values of *α* implying a more strict selection of best models (hence the choice of only the most performing features), the *SWAG* then uses the features selected in the first step to progressively build higher-dimensional models (i.e. models with an increased number of feature combinations within them) until it reaches the maximum number *p*_*max*_. When building the models for a given dimension, the *SWAG* takes the best models from the previous step (i.e. the step that built models with one less feature than the current step) and randomly adds a distinct feature from the set of features selected at the first step. Having built *m* models at each step (where *m* is also chosen by the user), the final output of the *SWAG* is a set of “strong” models (i.e. models with high predictive power) where each is based on a combination of 1 to *p*_*max*_ features. A simplified representation of the *SWAG* is presented in Fig 1. With this output, it is then possible for the user to apply post-processing to select a subset of interest from this set of models. The *SWAG* is made available as an `R` package on the CRAN repository. At the same time, a development version can also be found at https://github.com/SMAC-Group/SWAG-R-Package/.

**Fig 1.**
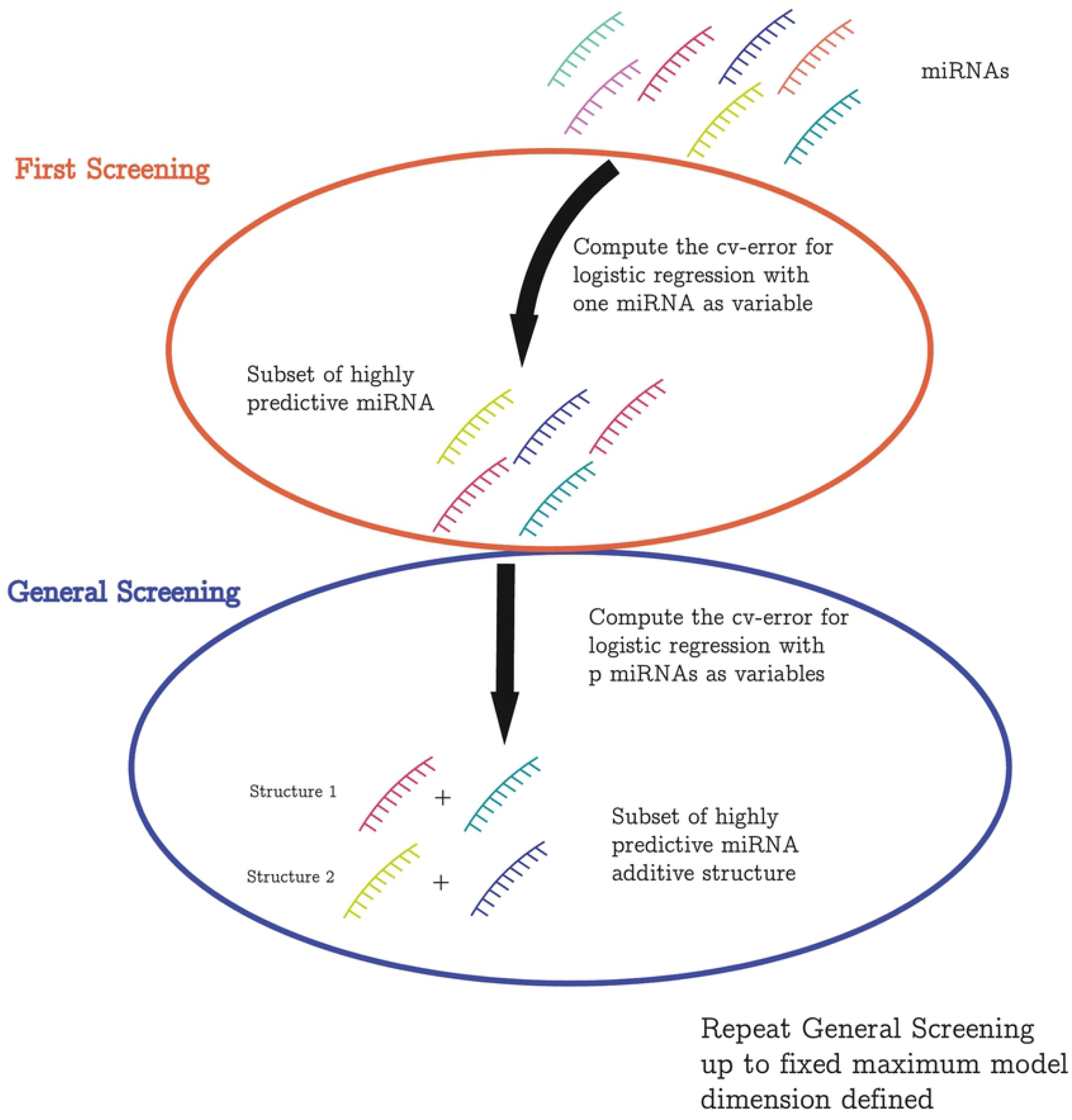
*SWAG* flowchart. A schematic representation of how the *SWAG* was calibrated for this work. The red set represents the first step which evaluates every one-dimensional model and selects the best expressions to be used in the general step represented by the blue set. The latter step evaluates and selects the best models of dimension 2 to *p*_max_.

### Implementation

The AHUS dataset is split into training and test subsets. The training subset contains 100 observations with a 56/44 split (normal tissue/invasive BC). The test subset has 25 observations with a 14/11 split. With CV, we estimate how accurately a predictive model will perform in practice. This is a well-established model validation technique for assessing how the results of a statistical analysis will generalize to an independent dataset. The aim is to estimate how accurately a model will perform in terms of prediction and the rationale of using this technique may be found in Fushiki [39] and Molinaro [10]. The caveats of CV are well explained in Bernau *et al*. [40] where the main setback can eventually consist in an overestimation of model performance in a broader application context.

Our research question is not about the validity of a given model selection method, but about the function of a specific miRNA in BC oncogenesis. To address this issue, we must not only select a set of models in which this specific miRNA is involved, but also determine the direction in which it acts (oncogenic or protective effect). Prior to this analysis, we have standardized the design matrix (miRNAs) to ensure a meaningful comparison across the different models. Then, to assess the role of the relevant miRNAs selected via the *SWAG*, we computed their *β* coefficients by performing a logistic regression on each element of the set of models. The evaluation of the *β* coefficients allow us to (i) identify either the oncogenic or protective effect of the variable and (ii) gain insight on its distribution. Positive values of *β* mean that the miRNA associated with this coefficient has an activating effect on tumor progression (oncogenic effect); a negative value means the opposite (protective effect). Since the miRNAs can be included in different models delivered by the *SWAG*, one can compute an empirical distribution of the coefficients.

### Single and associative effects on a binary variable

One of our research questions is whether a given miRNA has the same action (oncogenic or protective) when it is taken in isolation or when embedded within different models and feature combinations. In order to assess the biological action of the selected miRNAs, we compute both the single and associative effects of the *β* coefficients for each of the selected miRNAs. A single effect is measured by the estimated value of a *β* coefficient when considering a single miRNA in the logistic model. The associative effect is defined as all the different values that a *β* coefficient takes when constructing the set of models with the *SWAG*. The associative effect of a specific miRNA therefore may be seen as an indicator of its biological impact in a broader context.

As a matter of fact, according to Cox [17], these effects are typically different. For a random variable considered both alone and conditionally on a confounding variable W, the single and associative inferences may have opposite signs by the so-called Yule–Simpson effect [16]. This effect, if observed, may be explained in two ways : (i) the existence of sub-populations or (ii) the influence of a finite set of latent classes W (such as biological functions) within the population under study. An example of this phenomenon with some mathematical explanations can be found in Cox [17] and Boehm *et al*. [38]. Splitting the population into defined subgroups may, to some extent, dodge the first pitfall (i.e. the existence of msub-populations). However BC heterogeneity is large and has been documented in terms of different histological subtypes, treatment sensitivity profiles, and clinical outcomes. Furthermore, the heterogeneous expression of the oestrogen receptor, progesterone receptor, and HER2 has been reported in different areas of the same tumour. Molecular profiling studies have confirmed that spatial and temporal intratumour heterogeneity of BCs exist at a level beyond common expectations [41]. Splitting BC populations into subtypes may then be a misleading precaution. The second pitfall (i.e. the existence of latent biological functions shared by many genomic features) is even more elusive. Having not a single but a paradigmatic set of predictive models may help get around this hurdle. We addressed this challenge (differentiating the effect of sub-populations from that of latent biological variables on single and associative coefficients) by mixing the 55 invasive carcinomas into one category. If single and associative coefficients retain the same sign throughout the 112 selected models, we can conclude with some confidence that the existence of sub-populations (BC sub-types) has no effect on the oncogenic or protective effect of the relevant miRNAs inside the AHUS dataset. On the contrary, if single and associative coefficients have opposite signs, then one can assume that the effect of the relevant miRNA differs according to its environment.

### Horizontal and vertical organizations

We make the heuristic hypothesis that the human genome as a whole and its sub-units (such as non-coding RNAs) can be interpreted as semiotic systems. To give meaning to the miRNA-based net-like structures that we build through our statistical analysis, we borrow the notions of syntagm and paradigm from structural semiotic analysis, inspired by de Saussure theory [43]. A simple and useful introduction to semiotics may be found in Chandler [77]. De Saussure emphasized that meaning (in our case, oncogenic or protective effect) arises from differences between signifiers (in our case, miRNAs). These differences are of two kinds: syntagmatic (concerning positioning within a model) and paradigmatic (concerning substitution within a given model). These two dimensions are often presented as axes, where the horizontal axis is the syntagmatic and the vertical axis is the paradigmatic. The plane of the syntagm is that of the combination of signifiers (i.e. selected miRNAs) within a statistical model whilst the plane of the paradigm is that of the selection of signifiers. Whilst syntagmatic relations are combination possibilities, paradigmatic relations are functional contrasts. The meaning of a signifier is determined by both its paradigmatic and syntagmatic relations. According to this conception, the set made of all the selected models may be seen as the set of syntagmatic “sentences” selected by the *SWAG*, and the set made of the selected miRNAs as the set of paradigmatic Omics features. In this study, the horizontal syntagmatic axis will be used to tackle the second research question (to check how the behavior of the miRNAs varies according to the specific combination with which it interacts). The vertical paradigmatic axis will be used to address the third research question (to search for interchangeable miRNAs in these predictive models and by doing so, to decipher the biological targets of these miRNAs).

## Results

### Breast cancer / normal tissue discrimination

When applying *SWAG* to the AHUS data, a total of 45 miRNAs were selected, making a set of 112 indistinguishable models of sizes 4 to 5. These 112 models and the 45 selected miRNAs are displayed in the supporting information material. They perform similarly or outperform the lasso, a standard model selection method used in genomics [42], with less than half the number of miRNAs selected by the latter. This is evident from the comparison in terms of accuracy, sensitivity, specificity as well as positive and negative predictive values at the standard logistic cut-off level of 0.5 (cf. Fig 2). Indeed all the 112 *SWAG* models have an equal or greater accuracy than the lasso. Moreover a large majority of *SWAG* models (i.e. 97 out of 112) dominate lasso in any of the considered metrics. This can be inferred also visually in Fig 2 by looking at the barplots on the right of each specific *SWAG* range (i.e. the interval between the smallest and largest values among all SWAG models), with the vertical red line representing the corresponding lasso value. As a whole, these results suggest that *SWAG* more precisely targets the set of miRNAs involved in BC progression. Furthermore, Tab 1 reports two specific *SWAG* models, among the 112 selected ones, that achieve a perfect out-of-sample classification in terms of area under the curve (AUC). It is therefore possible to discriminate BC from normal breast tissue with extreme accuracy using miRNAs as biomarkers. The added value of the *SWAG* compared to lasso is that (i) it produces a set of equivalent models instead of a single one and (ii) the number of selected variables per model is smaller by a factor of two, making the models more easily interpretable.

**Table 1.** Best out-of-sample *SWAG* models that achieve a perfect classification in terms of area under the curve (i.e. *AUC =* 1).

| Non-coding miRNAs |  |  |  |  |  |
| --- | --- | --- | --- | --- | --- |
| Model 1 | hsa-miR-1274a | hsa-miR-21 | **hsa-miR-92a** | hsa-miR-328 | hsa-miR-140-3p |
| Model 2 | hsa-miR-1274a | hsa-miR-21 | **hsa-miR-92a** | hsa-miR-328 | **hsa-miR-30b** |

The non-coding miRNAs are displayed in order of presence in the SWAG chain. The chameleon miRNAs are presented in bold.

**Fig 2.**
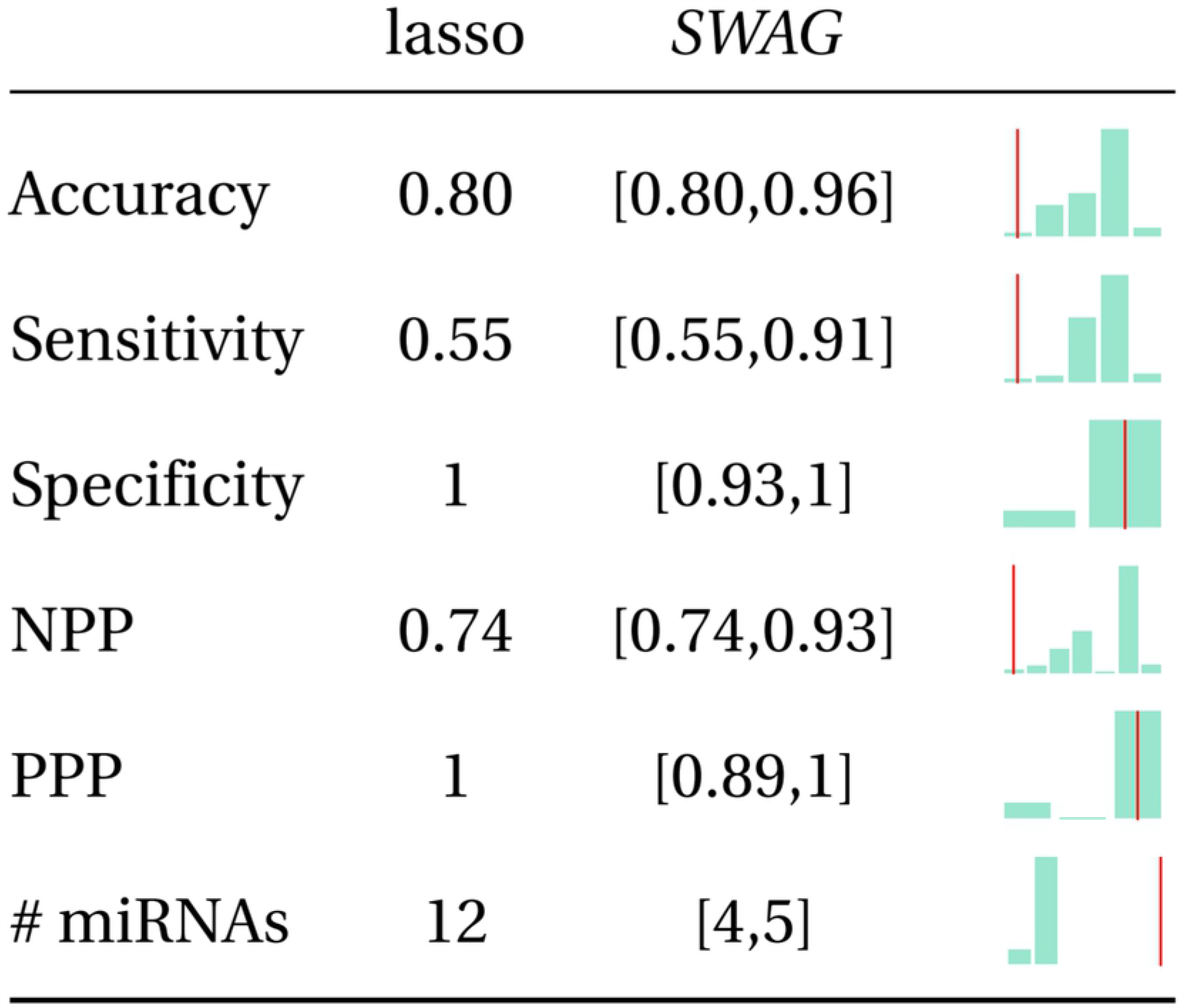
Comparison between Lasso and *SWAG*. We compare accuracy, sensitivity, specificity, negative predictive value (NPP), positive predictive value (PPP), number (#) of miRNAs of the lasso with the ranges (i.e. smallest-to-largest value intervals) of the same metrics for the 112 *SWAG* models. On the right of each *SWAG* range, a barplot illustrates the distribution of the specific metrics for the 112 considered models with a vertical red line representing the corresponding value for the lasso. All evaluations have been made at the standard 0.5 cut-off of logistic regression.

Among the 45 selected miRNAs, 8 were present in more than 16 % of all SWAG models. These 8 miRNAs are displayed in Tab 2, with their respective pairwise Spearman correlations, illustrated in Fig 3. As a non-parametric measure of rank correlation, Spearman correlation assesses how well the relationship between two variables can be described using a monotonic function. It is used in our study as an index of result consistency. Two miRNAs having similar effects on cancer progression should be positively correlated. The rational of this statement is that correlation and mutual information are closely related [76]. Selected miRNAs in each of the 112 statistical models are presented with single and associative *β* coefficients which are in most cases either positive (oncogenic effect) or negative (protective effect on tumor progression). However, one of them stands out from the others, `hsa-miR-92a`, whose associative *β* is positive and whose single one is negative (cf. Tab 3). This point will be discussed later.

**Table 2.** Model occurrence rate of the most frequently selected miRNAs.

| Non-coding miRNA | Model occurrence rate (%) |
| --- | --- |
| hsa-miR-1274a | 75.0 |
| hsa-miR-21 | 74.1 |
| hsa-miR-139-3p | 44.6 |
| hsa-miR-125b-2* | 39.3 |
| hsa-miR-92a | 25.0 |
| hsa-miR-449a | 22.3 |
| hsa-miR-155 | 18.8 |
| hsa-miR-200c | 16.1 |

**Table 3.**
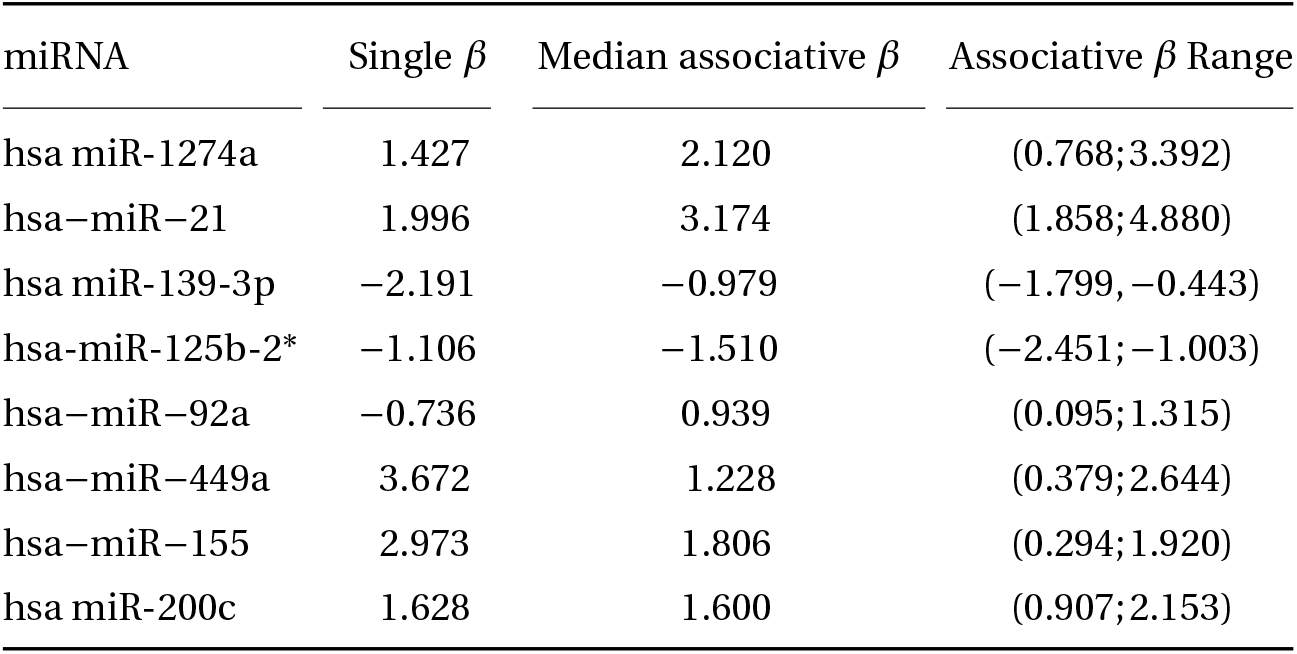
Single and associative coefficients (median values and range) for the eight most frequently selected miRNAs.

**Fig 3.**
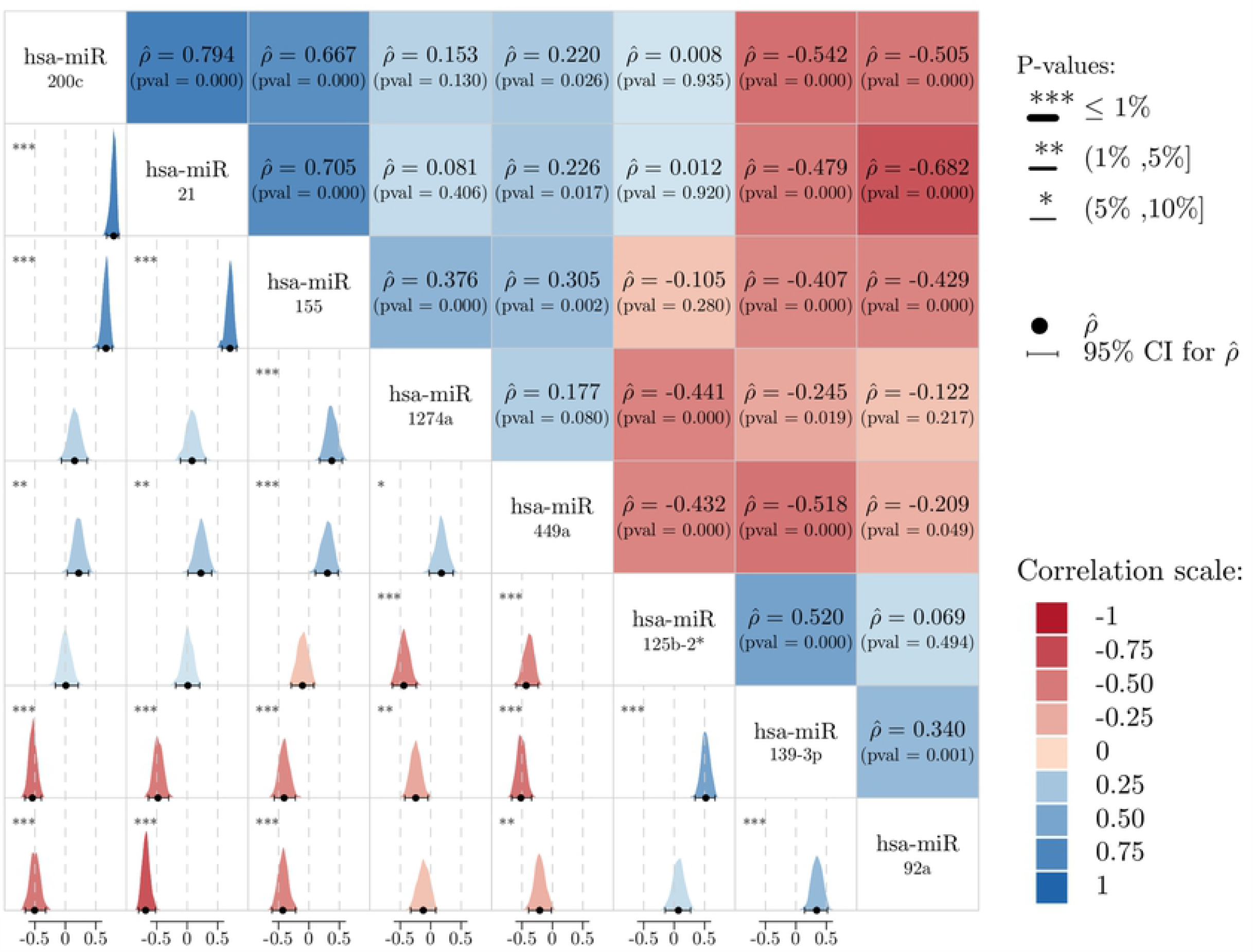
Spearman correlation 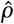 matrix for the 8 most frequent miRNAs selected by the *SWAG*. The upper triangular part shows the estimator 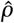 between miRNAs and their respective p-value computed via non-parametric bootstrap. The color of the boxes indicates the direction of 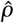 (blue for positive correlation and red for negative). The lower triangular part illustrates the bootstrap distribution of 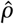 via the density plot, with the dark circle being the estimator of 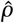 and the horizontal black line its 95% confidence interval. The star on the boxes’ upper-left indicates the level for which the correlation in significant.

### Oncogenic or protective role of miRNAs : the syntagmatic axis

The values of the single and associative coefficients of the eight most frequently selected miRNAs are displayed in Fig 4 and Tab 3. Coefficients related to miRNAs that are present in at least 10% of models and that show discordant values between single and associative *β*s are presented in Tab 4. Among the eight most frequently selected miRNAs, the median value of the *β*s considered along with with their range makes it possible to identify three classes of miRNA (cf. Fig 4 and Tab 3): (i) oncogenic miRNAs with single and associative positive values of *β* (i.e. `hsa-miR-1274a, hsa-miR-21, hsa-miR-449a, hsa-miR-155,` `hsa-miR-200c`); (ii) protective miRNAs (`hsa-miR-139-3p, hsa-miR-125b-2*`); (iii) an undefined miRNA (`hsa miR-92a`) with a positive single coefficient and a negative associative one. An indication of the consistency of these results lies in the Spearmann correlation coefficients 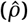 (cf. Fig 3): `hsa-miR-1274a` is significantly positively correlated (at the level *α =* 5%) to its oncogenic partner `hsa-miR-155` and negatively correlated to protective miRNAs `hsa-miR-125b-2*` and `hsa-miR-139-3p`. The same consistency can be observed for `hsa-miR-21`. The two protective miRNAs, `hsa-miR-139-3p` and `hsa-miR-125b-2*`, are also significantly positively correlated with each other, and significantly negatively correlated to oncogenic miRNAs such as `hsa-miR-449a` and `hsa-miR-1274a`.

**Table 4.** Single and associative coefficients (median values and range) for the *“chameleon”* miRNAs present in at least 10% of the models.

| miRNA | Single $\beta$ | Median associative $\beta$ | Associative $\beta$ Range |
| --- | --- | --- | --- |
| hsa-miR-92a | -0.736 | 0.939 | (0.095;1.315) |
| hsa-miR-320d | -1.064 | 1.174 | (-0.412;2.543) |
| hsa-miR-193a-5p | -1.734 | 1.114 | (-1.318,1.225) |
| hsa-miR-30b | 0.694 | -0.753 | (-1.422;0.595) |

Associative ranges and not confidence intervals are shown since some of the coefficients display a bi-modal distribution.

**Fig 4.**
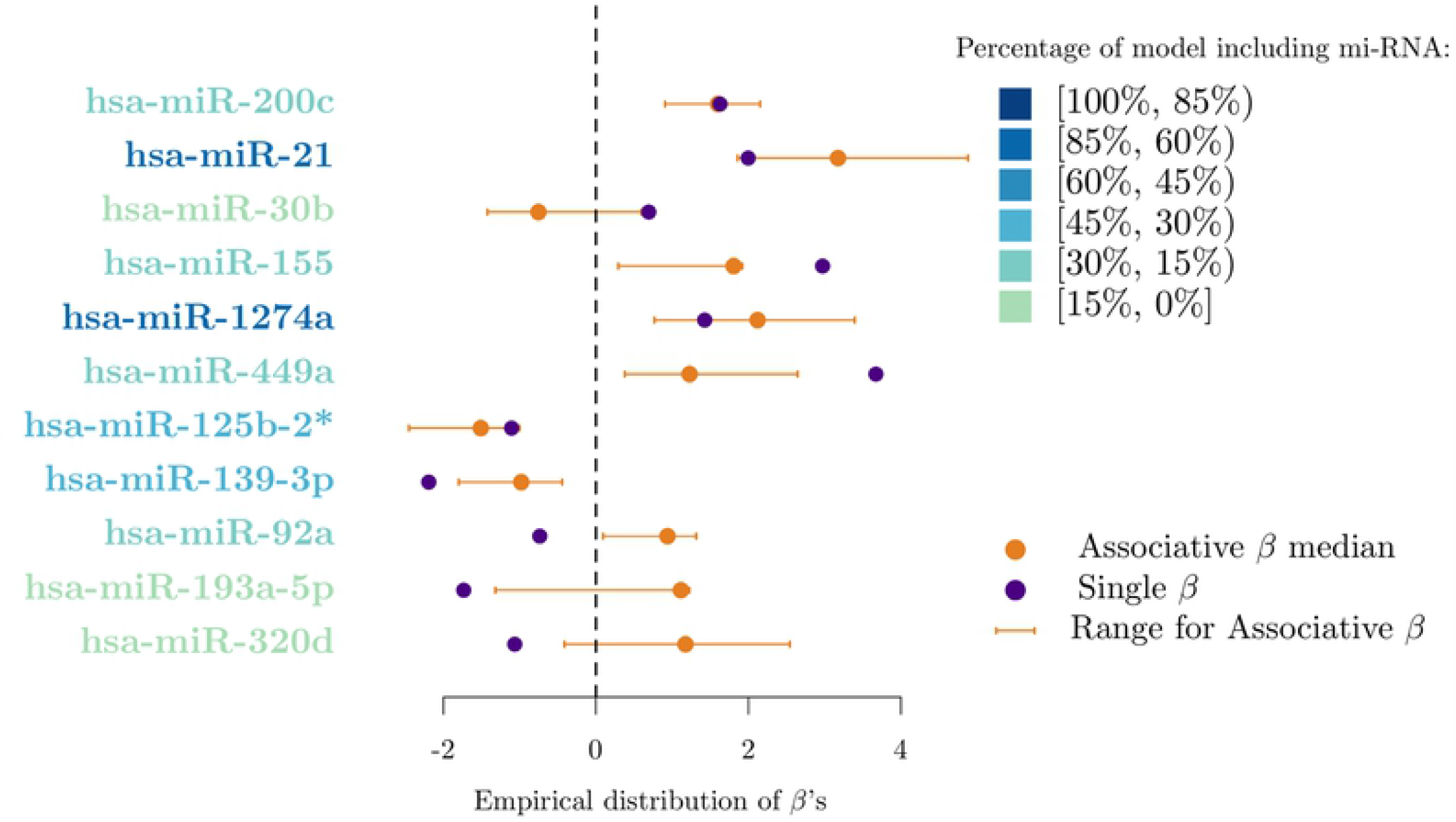
Distribution of *β* coefficients for the most frequently selected miRNAs. We present the single effect (i.e. the estimated value of a *β* coefficient when considering a single miRNA in the logistic model), the median and range of the associative effect (i.e. the different values that a miRNA specific *β* coefficient takes in each of the 112 *SWAG* models) for each of the most frequently selected miRNAs displayed in both Tab 3 and Tab 4.

The case of `hsa-miR-92a` which displays discordant single and associative coefficients is not isolated. Among the miRNAs selected in at least 10% of the models, three show a similar behaviour to that of `hsa-miR-92a`, `hsa-miR-320d`, `hsa-miR-193a-5p` and `hsa-miR-30b`. Using a bioinformatics-based interaction analysis of `hsa-miR-92a-3p` and key genes in tamoxifen-resistant BC cells, Cun *et al*. [69] found that `hsa-miR-92a-3p` was higher in BC serum or tissue than in healthy volunteer serum or adjacent normal tissue. Hence, high expression of `hsa-miR-92a-3p` seems to predict poor prognosis for BC patients according to this meta-analysis study. This finding is in contradiction with previous results published by Nilsson *et al*. [70] that suggest that downregulation of `hsa-miR-92a-3p` is associated with aggressive BC features and increased tumour macrophage infiltration. In relation to `hsa-miR-320d` action in BC, Cava *et al*. [86] found that its downregulation favours BC progression. To the best of our knowledge, no other study has investigated the role of `hsa-miR-320d` in BC progression therefore it is not possible to compare our result with other data coming from recent literature. According to Maltseva *et al*. [87], `hsa-miR-193a-5p` is less expressed in inflammatory BC patients and is known to play a suppressive role in BC. This statement is in contradiction with the findings in Li *et al*. [88] that state that long non-coding RNA small nucleolar RNA host gene 1 (SNG1) activates HOXA1 expression via sponging `hsa-miR-193a-5p` in BC progression. Finally, the role of `hsa-miR-30b` has been shown to be versatile, as a recent review points out [89]. Members of the `hsa-miR-30` family play a role in the regulation of tumorigenesis, interference with tumor invasion and metastasis as well as reversal of drug resistance. Nevertheless some `hsa-miR-30` family members have independent protective effects on the prognosis of BC patients. In conclusion, a versatile role of miRNAs in BC progression is quite a common finding in recent literature. Our results give a statistical basis to this allegation.

### An example of paradigmatic substitution: hsa-miR-140-3p and hsa-miR-30b

Among the 112 selected models, two of them show a perfect discriminating power (cf. Tab 1). These two differ only by one miRNA: `hsa-miR-140-3p` and `hsa-miR-30b` respectively. From a linguistic point of view, these two miRNAs can be seen as synonyms, meaning that they can be swapped without affecting the meaning of the “sentence” (the predictive power of the model). As stated previously, miRNAs are short endogenous noncoding RNAs that regulate their target messenger RNAs by promoting their degradation or by repressing their translation [44]. It is therefore intuitive to look at the set of target mRNAs associated with each of these regulatory factors and to determine their intersection. By doing so, we can create a list of common target genes. We acquired information on the targets of these two miRNAs from the http://mirbase.org/ platform [90], [91], [92], [93], [94], and [95]. The targets of these two miRNAs were obtained by crossing information from the TargetScanvert database (http://www.targetscan.org/) [83], [85] and the miRDB database (http://mirdb.org/) [84]. The results are shown in Tab 5.

**Table 5.** hsa-miR-140-3p and hsa-miR-30b common targets.

| Gene symbol | Gene name |
| --- | --- |
| NDST1 | N-deacetylase and N-sulfotransferase 1 |
| CCNT2 | Cyclin T2 |
| USP49 | Ubiquitin specific peptidase 49 |
| ZC3H6 | Zinc finger CCCH-type containing 6 |
| DTX4 | Deltex E3 ubiquitin ligase 4 |
| CDK6 | Cyclin dependent kinase 6 |
| SRGAP3 | SLIT-ROBO Rho GTPase activating protein 3 |
| RAB21 | RAB21, member RAS oncogene family |
| P2RY2 | Purinergic receptor P2Y2 |
| GLG1 | Golgi glycoprotein 1 |
| KCNB1 | Potassium voltage-gated channel subfamily B member 1 |
| TNKS | Tankyrase |
| UBN2 | Ubinuclein 2 |
| RFT1 | RFT1 homolog |
| ARID2 | AT-rich interaction domain 2 |
| TYRO3 | TYRO3 protein tyrosine kinase |
| EIF5A2 | Eukaryotic translation initiation factor 5A2 |
| HPCAL4 | Hippocalcin like 4 |

These targets were selected by crossing information coming from the http://mirdb.org/ and http://www.targetscan.org/ databases

Among these target genes common to `hsa-miR-140-3p` and `hsa-miR-30b`, some are well known to play a pivotal role in cancer progression. In this direction, one can cite USP49 [96], CDK6 [97], RAB21 [98], P2RY2 [99], TNKS [100], ARID2 [101], TYRO3 [102], EIF5A2 [103]. This set of common targets may explain why these two miRNAs are “synonyms” and can be exchanged in predictive models without any harm. The latent functions of these putative target genes is shown in Tab 6.

**Table 6.** Function of hsa-miR-140-3p and hsa-miR-30b targets involved in cancer pathophysiology.

| Gene symbol | Gene Function |
| --- | --- |
| USP49 | Histone H2B lysine deubiquitination / mRNA splicing |
| CDK6 | Cell dedifferentiation / cell division |
| RAB21 | Rab protein signal transduction |
| P2RY2 | Cellular ion homeostasis / cellular response to ATP |
| TNKS | Cell division / mitotic spindle organisation |
| ARID2 | Negative regulation of cell migration and cell population proliferation |
| TYRO3 | Apoptotic cell clearance / cell adhesion and migration |
| EIF5A2 | mRNA transport / regulation of cell population proliferation |

Source : https://ensembl.org/ and https://uniprot.org/

## Discussion

With regards to our research questions, we can firstly conclude that it is possible to differentiate normal breast tissue from breast carcinoma by using miRNAs as biomarkers with reasonable sensitivity and specificity. Secondly, some selected miRNAs behave in opposite ways according to the models in which they are embedded. We decided to call these miRNAs, whose action is conditioned by their insertion in a syntagmatic axis, *chameleon* micro RNAs. Thirdly, any model selection method such as the one used for this work (*SWAG*) that gives the opportunity to build “horizontal” and “vertical” axes can point to latent biological functions and help researchers develop new hypotheses. In our case, with regards to `hsa-miR-140-3p` and `hsa-miR-30b`, some latent cell functions such as cell division and differentiation, mRNA splicing and transport as well as cellular ion homeostasis appear to be highly relevant.

According to Stepanenko *et al*. [29], cancer evolution is a stochastic process both at the genome and gene levels. Most tumors contain multiple genetic subclones, evolving in either succession or in parallel, either in a linear or branching manner, with heterogeneous genome and gene alterations, extensively rewired signaling networks, and addicted to multiple oncogenes easily switching with each other during cancer progression and medical intervention. Hundreds of discovered cancer genes or gene products are classified according to whether they function in an oncogenic or protective manner in a cancer cell. However, there are many cancer “*gene-chameleons*”, which behave in opposite manners in different experimental settings showing what Stepanenko calls “antagonistic duality”. These statements find confirmation in our study. This antagonistic duality affects not only genes, but also miRNAs. For this subgroup, the distribution of the *β* coefficients, either single or associative, include the value zero thereby indicating an ambiguous or dualistic effect. These results are in line with the most recent literature about their action in BC progression. Further research taking into account this versatile effect according to net-like structures is needed. Our research places itself within the emerging field of artificial intelligence [11]. With the advent of Big Data and the ever-increasing storage and computing power, the challenge has shifted from collecting data to turning it into meaningful and actionable insights. This challenge requires that we leave on the side of the road statistical methods that select genomic items taken in isolation and that we favour methods that scrutinize biological systems. By producing net-like combinations of equivalent models, it is possible to shed light on the latent biological confounding variables that are usually ignored and may reverse the effect of the considered Omics feature.

## Added value

The added value of our research is fourfold:

- Predictive models with indistinguishable (optimal) predictive abilities are not unique, but belong to a set of equivalent and, in some sense, exchangeable models.
- Our results indicate that miRNAs are not isolated items but are integrated into two-dimension statistical axes. Their function cannot be inferred independently of the other components of the syntagmatic or horizontal axis.
- Some miRNAs are exchangeable in terms of predictive ability and point to latent biological functions.
- Conflicting results in the literature suggest that a protective or an oncogenic effect cannot be definitely assigned to any miRNA. Data-driven nets may help biologists in building new hypotheses and experimental designs in order to decipher the function of non-coding RNA, which may act as *chameleon* molecules according to the organization in which they are embedded.

## Supporting Information

Below we state all the material that is auxiliary to the main content of the article and can be found in a separate file.

- **S1 Table 1. List of all the 45 miRNAs selected by the *SWAG***.
- **S1 Table 2. List of all the models with 4 miRNAs selected by the *SWAG***.
- **S1 Table 3. List of all the models with 5 miRNAs selected by the *SWAG*. (Part 1)**
- **S1 Table 4. List of all the models with 5 miRNAs selected by the *SWAG*. (Part 2)**
- **S1 Table 5. List of all the models with 5 miRNAs selected by the *SWAG*. (Part 3)**

## Acknowledgments

We thank Haakensen *et al*. for having made available the AHUS data set on the free access ArrayExpress platform at: https://www.ebi.ac.uk/arrayexpress/experiments/E-MTAB-3759/?query=AHUS. The statistical analysis performed in this study is based on their data.

## Conflict of Interest Statement

### Ethics approval and consent to participate

Not applicable.

### Consent for publication

Not applicable.

### Competing interests

The authors declare that they have no competing interests.

